# The microbiome and metabolome of pre-term infant stool is personalized, and not driven by health outcomes including necrotizing enterocolitis and late-onset sepsis

**DOI:** 10.1101/125922

**Authors:** Stephen Wandro, Stephanie Osborne, Claudia Enriquez, Christine Bixby, Antonio Arrieta, Katrine Whiteson

## Abstract

The assembly and development of the gut microbiome in infants has important consequences for immediate and long-term health. Preterm infants represent an abnormal case for bacterial colonization because of early exposure to bacteria and frequent use of antibiotics. To better understand the assembly of the gut microbiota in preterm infants, fecal samples were collected from 32 very low birthweight preterm infants over the first six weeks of life. Infant health outcomes included healthy, late-onset sepsis, and necrotizing enterocolitis (NEC). We characterized the bacterial composition by 16S rRNA gene sequencing and metabolome by untargeted gas chromatography mass spectrometry. Preterm infant fecal samples lacked beneficial *Bifidobacterium* and were dominated by *Enterobacteriaceae, Enterococcus*, and *Staphylococcus* due to the near uniform antibiotic administration. Most of the variance between the microbial community compositions could be attributed to which baby the sample came from (Permanova R^2^=0.48, p<0.001), while clinical status (healthy, NEC, or late-onset sepsis), and overlapping time in the NICU did not explain a significant amount of variation in bacterial composition. Fecal metabolomes were also found to be unique to the individual (Permanova R^2^=0.43, p<0.001) and weakly associated with bacterial composition (Mantel statistic r = 0.23 ± 0.05 (p = 0.03 ± 0.03). No measured metabolites were found to be associated with necrotizing enterocolitis, late-onset sepsis or a healthy outcome. Overall, preterm infants gut microbial communities were personalized and reflected antibiotic usage.

**Importance:** Preterm infants face health problems likely related to microbial exposures including sepsis and necrotizing enterocolitis. However, the role of the gut microbiome in preterm infant health is poorly understood. Microbial colonization differs from healthy term babies because it occurs in the NICU and is often perturbed by antibiotics. We measured bacterial compositions and metabolomic profiles of 77 fecal samples from thirty-two preterm infants to investigate the differences between microbiomes in health and disease. Rather than finding microbial signatures of disease, we found the preterm infant microbiome and metabolome were both personalized, and that the preterm infant gut microbiome is enriched in microbes that commonly dominate in the presence of antibiotics. These results contribute to the growing knowledge of the preterm infant microbiome and emphasize that a personalized view will be important to disentangling the health consequences of the preterm infant microbiome.

## Introduction

Early life exposure to microbes and their metabolic products is a normal part of development, with enormous and under-explored impact on the immune system. The intestinal microbiota of infants initially assembles by exposure to the mother’s microbiota and microbes in the environment (1–4). In healthy breast-fed infants *Bifidobacteria longum* spp. infantis capable of digesting human-milk oligosaccharides dominate the infant gut (5). When infants are born preterm, they are exposed to environmental and human associated microbes earlier in their development than normal, and rarely harbor *Bifidobacteria* spp. in their gut communities. We do not yet understand the effects of altering the timing of initial bacterial exposure. With numerous emerging health consequences related to the microbiome, understanding factors that influence this initial assembly of the microbiome will be important.

Preterm infants are routinely treated with antibiotics, enriching for microbes that can colonize in the presence of antibiotics (4, 6, 7). While antibiotics have tremendously reduced infant mortality, their effect on microbiota assembly and resulting health consequences is not fully understood. Prenatal and postnatal antibiotics have been shown to reduce the diversity of the infant intestinal microbiota (8, 9). Children under two years old are prescribed antibiotics at a higher rate than any other age group, and 85% of extremely low birthweight infants (< 1000 g) are given at least one course of antibiotics (10). Even if an infant is not exposed to antibiotics after birth, approximately 37% of pregnant women use antibiotics over the course of the pregnancy (11).

Perturbing the microbiota of infants can have immediate and long-lasting health consequences. In the short term, infants can be infected by pathogenic bacteria that results in sepsis, which is categorized as early-onset or late-onset depending on the timing after birth. Preterm infants are also at high risk to develop necrotizing enterocolitis (NEC), which is a devastating disease that causes portions of the bowel to undergo necrosis. NEC is one of the leading causes of mortality in preterm infants, who make up 90% of NEC cases (12). The incidence of NEC among low birthweight preterm infants is approximately 7% and causes death in about one third of cases. The exact causes of NEC are not known, but an excessive inflammatory response to intestinal bacteria may be involved (13).

Many of the long-term consequences of microbial colonization are believed to be mediated by interactions between the intestinal microbiota and the immune system. In addition to direct interactions, the microbiota interacts with the immune system through the production of metabolites that can be taken up directly by immune and epithelial cells (14, 15). For example, bacterial production of short chain fatty acids can affect health and integrity of the intestinal epithelia and immune cells (16–18). However, few studies use metabolites alongside bacterial community profiling. In fact, the healthy composition of an infant fecal metabolome is understudied.

In this retrospective study, we follow the changes in the gut microbiota over time in 32 very low birth weight (< 1500 grams) preterm infants born at Children’s Hospital Orange County. We simultaneously track the bacterial composition and metabolite profile over time. Infants were classified into three groups based on health outcomes: healthy infants, late-onset sepsis, and NEC. The composition of the intestinal microbiota was measured by 16S rRNA gene sequencing of fecal samples taken over time. Preterm infant guts were dominated by *Enterobacteriaceae* and *Enterococcus*, and *Staphylococcus.* Untargeted metabolomics analysis of the fecal samples by gas chromatography mass spectrometry (GC-MS) revealed a personalized metabolome that was weakly associated with the bacterial composition.

## Results

### Patient cohort

A total of 77 fecal samples were collected from 32 very low birth weight infants in the NICU at Children’s Hospital Orange County from 2011 to 2014 (Table 1, Figure 1). Birthweights ranged from 620 – 1570 grams. Fecal samples were collected between day 7 and 75 of life. Sampling time and number of fecal samples varied. Three or more longitudinal samples were available from ten of the infants, while one or two samples were available from the remaining 22 infants. Three infants developed NEC, eight developed late-onset sepsis, and 21 remained healthy. Twelve infants were delivered vaginally while the remaining 22 were delivered by cesarean section. All infants were fed by either breastmilk or a combination of breastmilk and formula. Twenty-four infants received antibiotics at some point during the sampling period, the most common being ampicillin and gentamycin.

**Table 1.** Clinical and sampling information for all infants.

| Infant | # samples | Age sample(s) collected (days) | Age at disease onset (days) | Group | Fetal age at birth | Birth weight (g) | Antibiotics | Delivery mode | Feeding type | Twin set |
| --- | --- | --- | --- | --- | --- | --- | --- | --- | --- | --- |
| 1 | 2 | 7,7 |  | control | 27w4d | 875 |  | c-section | breastmilk |  |
| 2 | 3 | 15,15,36 |  | control | 31w | 1570 | ampicillin,gentamycin | c-section | breastmilk and formula |  |
| 3 | 1 | 19 |  | control | 26w | 980 | ampicillin,gentamycin | c-section | breastmilk |  |
| 4 | 2 | 11,11 |  | control | 30w3d | 1335 |  | vaginal | breastmilk |  |
| 5 | 2 | 18,18 |  | control | 24w5d | 630 |  | c-section | breastmilk |  |
| 6 | 4 | 25,26,28,43 |  | control | 28w5d | 860 | ampicillin,gentamycin | c-section | breastmilk and formula |  |
| 7 | 3 | 10,21,24 |  | control | 25w2d | 885 |  | c-section | breastmilk and formula |  |
| 8 | 1 | 10 |  | control | 25w4d | 940 | ampicillin,gentamycin | vaginal | breastmilk |  |
| 9 | 1 | 8 |  | control | 27w2d | 1205 |  | vaginal | breastmilk |  |
| 11 | 2 | 29,29 |  | control | 27w4d | 850 | ampicillin,gentamycin | vaginal | breastmilk |  |
| 12 | 1 | 22 |  | control | 26w2d | 880 | ampicillin,gentamycin | c-section | breastmilk and formula | 1 |
| 13 | 1 | 23 |  | control | 26w2d | 925 | ampicillin,gentamycin | c-section | breastmilk and formula | 1 |
| 14 | 1 | 8 |  | control | 31w4d | 1190 | ampicillin,gentamycin | c-section | breastmilk |  |
| 15 | 3 | 18,40,40 |  | control | 28w1d | 1270 | ampicillin,gentamycin | c-section | breastmilk and formula | 2 |
| 16 | 1 | 19 |  | control | 28w1d | 1355 | ampicillin,gentamycin | c-section | breastmilk and formula | 2 |
| 17 | 3 | 18,32,54 |  | control | 26w2d | 660 | ampicillin,cefotaxime | c-section | breastmilk |  |
| 21 | 1 | 10 |  | control | 28w6d | 1180 | ampicillin,gentamycin | c-section | breastmilk |  |
| 22 | 1 | 25 |  | control | 28w6d | 1360 | ampicillin,cefotaxime | vaginal | breastmilk and formula |  |
| 24 | 2 | 27,73 |  | control | 26w | 740 | ampicillin,gentamycin | c-section | breastmilk | 3 |
| 25 | 1 | 28 |  | control | 26w | 780 | ampicillin,gentamycin | c-section | breastmilk | 3 |
| 35 | 2 | 18,18 |  | control | 25w5d | 920 |  | c-section | breastmilk and formula |  |
| 23 | 7 | 14,15,27,28,30,30,56 | 27 | NEC | 26w6d | 1080 | ampicillin,gentamycin, cefotaxime,vancomycin | vaginal | breastmilk |  |
| 28 | 4 | 31,32,33,48 | 31 | NEC | 26w | 1060 | vancomycin,piperacillin | c-section | breastmilk and formula |  |
| 30 | 4 | 21,41,42,56 | 41 | NEC | 23w6d | 620 | cefazolin,azithromycin,ampicillin | vaginal | breastmilk and formula |  |
| 20 | 1 | 21 | 26 | septic | 24w5d | 815 | ampicillin,gentamycin | c-section | breastmilk |  |
| 10 | 6 | 15,35,36,37,39,40 | 27 | septic | 26w5d | 940 | ampicillin,gentamycin,vancomycin | vaginal | breastmilk and formula |  |
| 26 | 1 | 22 | 22 | septic | 24w4d | 660 | ampicillin,gentamycin,cefotaxime,vancomycin | c-section | breastmilk | 4 |
| 27 | 2 | 22,31 | 29 | septic | 24w5d | 650 | ampicillin,gentamycin | c-section | breastmilk | 4 |
| 29 | 2 | 20,26 | 26 | septic | 26w1d | 980 | cefotaxime,vancomycin | c-section | breastmilk |  |
| 31 | 5 | 10,34,35,38,45 | 34 | septic | 27w | 710 | ampicillin,gentamycin | c-section | breastmilk and formula |  |
| 32 | 4 | 32,32,53,75 | 32 | septic | 27w5d |  |  |  | breastmilk and formula |  |
| 37 | 3 | 8,17,18 | 13 | septic | 24w1d | 680 | ampicillin,gentamycin,cefazolin,oxacillin | vaginal | breastmilk |  |

**Figure 1.**
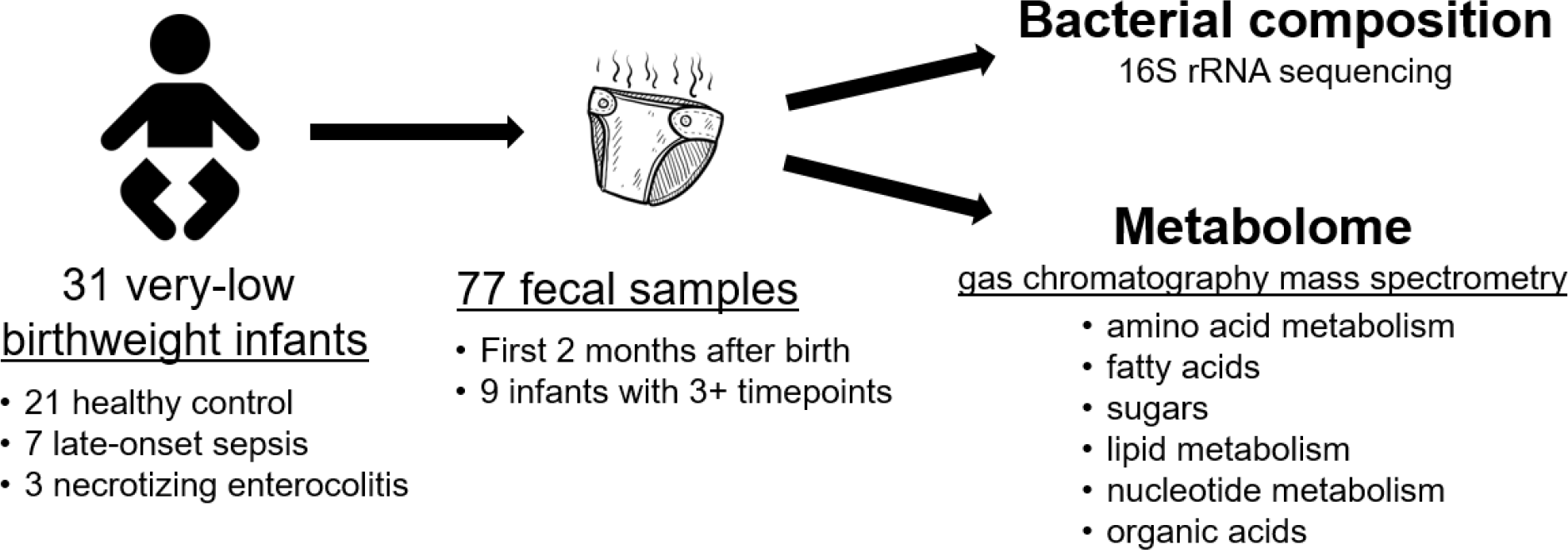
Study design schematic. Longitudinal fecal samples were collected over the first 75 days of life from very low birthweight infants in the NICU. Bacterial compositions and metabolomes were characterized.

### Microbial Community Characterization

We sequenced the 16S rRNA gene content of each fecal sample to determine bacterial composition. The total bacterial load of each fecal sample was measured by qPCR of the 16S rRNA gene and scaled to the total weight of stool that DNA was extracted from. Among all infants, bacterial abundances vary over four orders of magnitude and were not associated with health outcome (**Supplemental figure 1**). The high variation in bacterial load is likely due to the near uniform use of antibiotics. Bacterial communities were composed of mostly *Enterobacteriaceae, Enterococcus, Staphylococcus*, and *Bacteroides* (Figure 2a). Most samples were dominated by one to three genera of bacteria. Only three infants (two fed breastmilk, one fed breastmilk and formula) were colonized at greater than 1 % relative abundance by *Bifidobacteria*, which emerging evidence suggests is a key member of the infant microbiome. We computationally confirmed that the primers used are able to detect 90% of known *Bifidobacteria* species (19). No single bacterial OTU or community composition was consistently found for infants that became sick (NEC or late-onset sepsis) compared to the infants that remained healthy.

**Figure 2.**
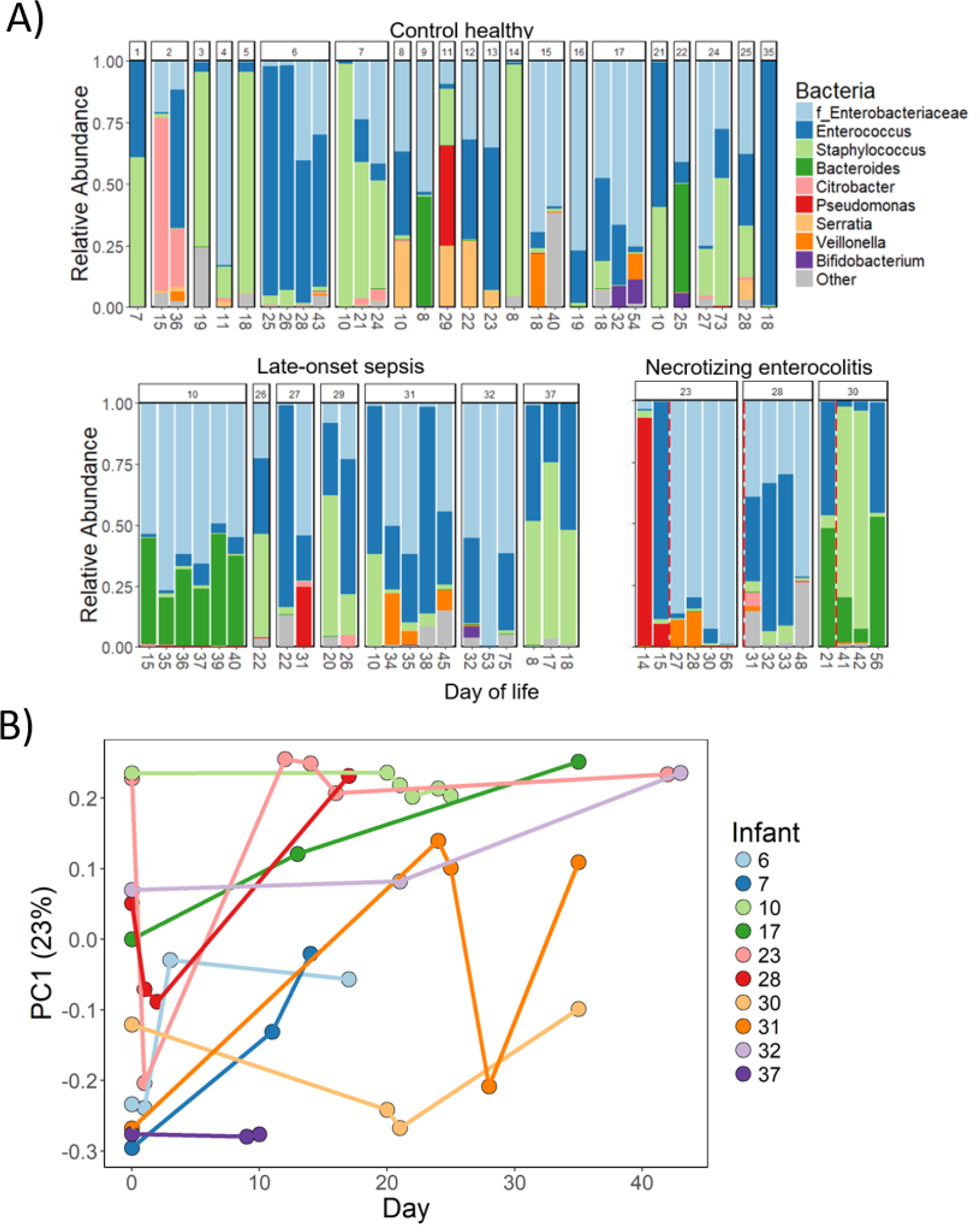
Bacterial composition of preterm infant guts. A) Stacked barplot of relative abundance of bacterial genera in all infant samples. The family *Enterobacteriaceae* is included because genus level resolution was not available. Infants are grouped together by health outcome. The timing of necrotizing enterocolitis diagnosis is indicated by a vertical dotted line. B) First axis of PCoA based on weighted uniFrac distances between bacterial communities plotted over time. Only infants with three or more longitudinal samples shown in B.

Longitudinal sampling revealed that over the course of days, the bacterial composition could change dramatically (Figure 2a, 2b). Permutational Multivariate Analysis of Variance (PERMANOVA) was applied to determine which of the known clinical factors explained the most variance in the bacterial community composition. The individual explained 48% (p < 0.001) of the variance in the samples, meaning that about half of the total variance among all tested fecal samples could be attributed to the infant the fecal sample came from (**Supplemental table 1**). None of the other factors explained a significant amount of variation in the bacterial composition, including infant health, overlapping dates in the NICU, delivery mode, or feeding mode. Four of the infants in the study are twins. Twin set 1 (infants 12 and 13) had a similar microbial composition while the other three sets did not (**Supplemental Figure 2**).

Diversity of the bacterial communities was low as expected for preterm infants. Alpha diversity as measured by Shannon index increased overall with age, but the trend was not significant (linear model R^2^ = 0.005, p = 0.52) (Figure 3a). Other clinical factors including health outcome, feeding (breastmilk versus breastmilk and formula), antibiotic use, and delivery mode were tested for an effect on the alpha diversity (Figure 3b-e). None of the factors were associated with a difference in alpha diversity except recorded antibiotic use, in which Shannon diversity was unexpectedly lower on average in infants that did not have a record of receiving antibiotics (Wilcoxon rank sum test p=0.06). It should be noted that although six infants did not have a record of antibiotic use, records may be incomplete due to hospital transfers. All four infants that were colonized with *Bacteroides* were born vaginally, although five other vaginally born infants were not colonized. Only vaginally born infants were colonized by *Bacteroides* (four out of nine infants) while none of the twenty-two infants born by C-section were colonized.

**Figure 3.**
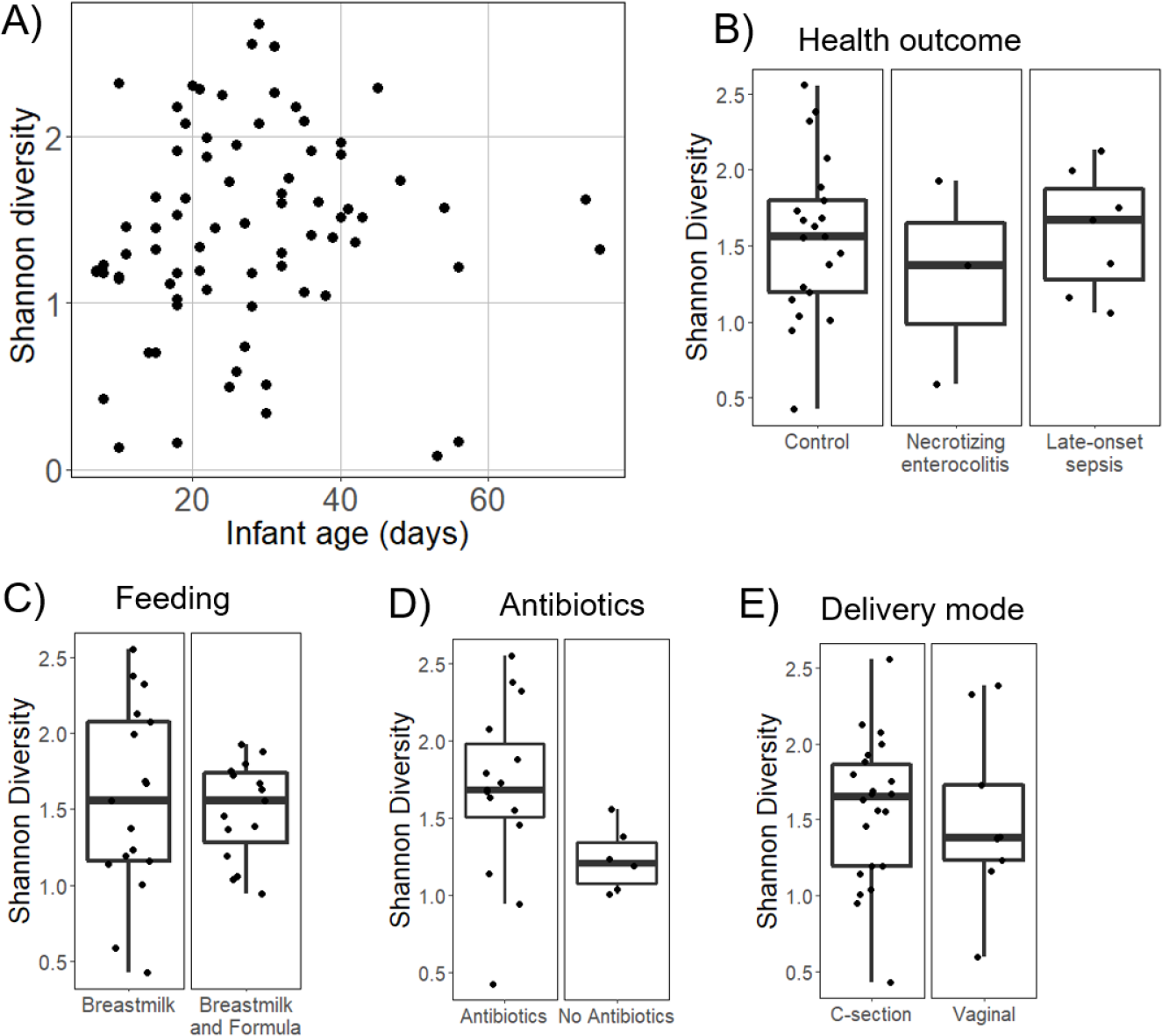
Alpha diversity measured by Shannon index of bacterial composition. A) Alpha diversity of all samples over the age of the infant. Boxplots of the average alpha diversity of each infant separated by B) health outcome, C) infants that were fed only breastmilk or a combination of formula and breastmilk, D) record of antibiotic usage, or E) delivery mode.

### Metabolomics

Metabolite profiles of infant fecal samples were analyzed by gas chromatography mass spectrometry, which measures small primary metabolites. Over 400 small molecules were detected from each fecal sample and 224 metabolites were known compounds. Metabolites were grouped into the following categories: amino acid metabolism, bile acids, central metabolism, fatty acids, fermentation products, lipid metabolism, nucleotide metabolism, organic acids, sterols, sugars, sugar acids, sugar alcohols, and vitamin metabolism (Figure 4, **Supplemental table 2**). No metabolites or categories of metabolites were found to be associated with necrotizing enterocolitis or late-onset sepsis. The metabolite profile of each infant was seen to vary over time, similar to the amount of variation seen in the bacterial composition (Figure 5). PERMANOVA analysis to determine which factors explain the most variance in the metabolite profile indicate that the individual explains 43% (p < 0.001) of the variation (**Supplemental table 1**).

**Figure 4.**
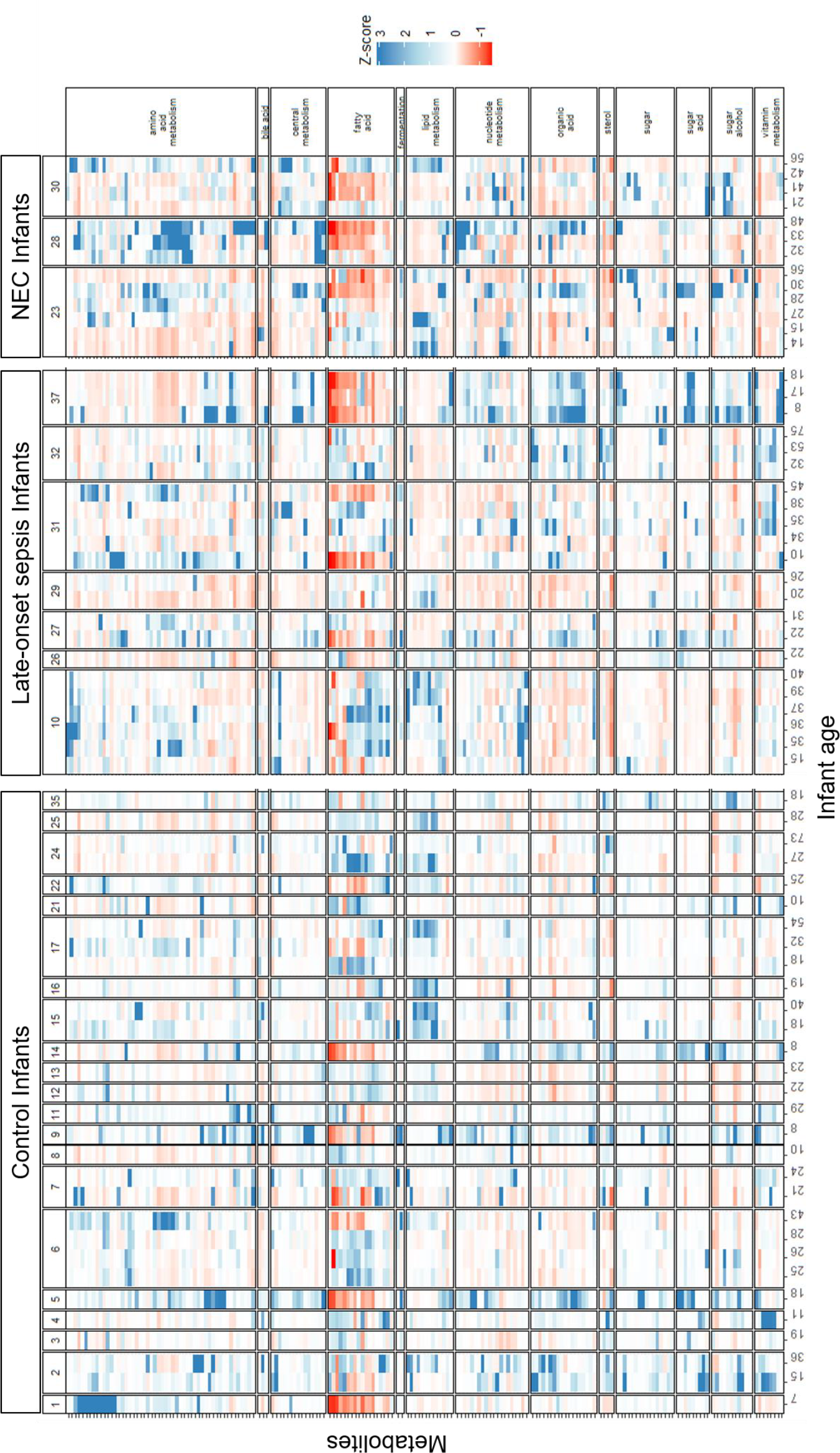
Metabolite profile of preterm infant fecal samples. Color is the modified z-score which is based on the median intensity for each metabolite in all infant samples. Red cells indicate standard deviations below the median and blue indicate standard deviations above the median value for each metabolite. Measured metabolites that could be assigned to a category are shown. Samples on the x-axis and grouped by infant and ordered longitudinally. Metabolites within each category are listed in the supplemental data.

**Figure 5.**
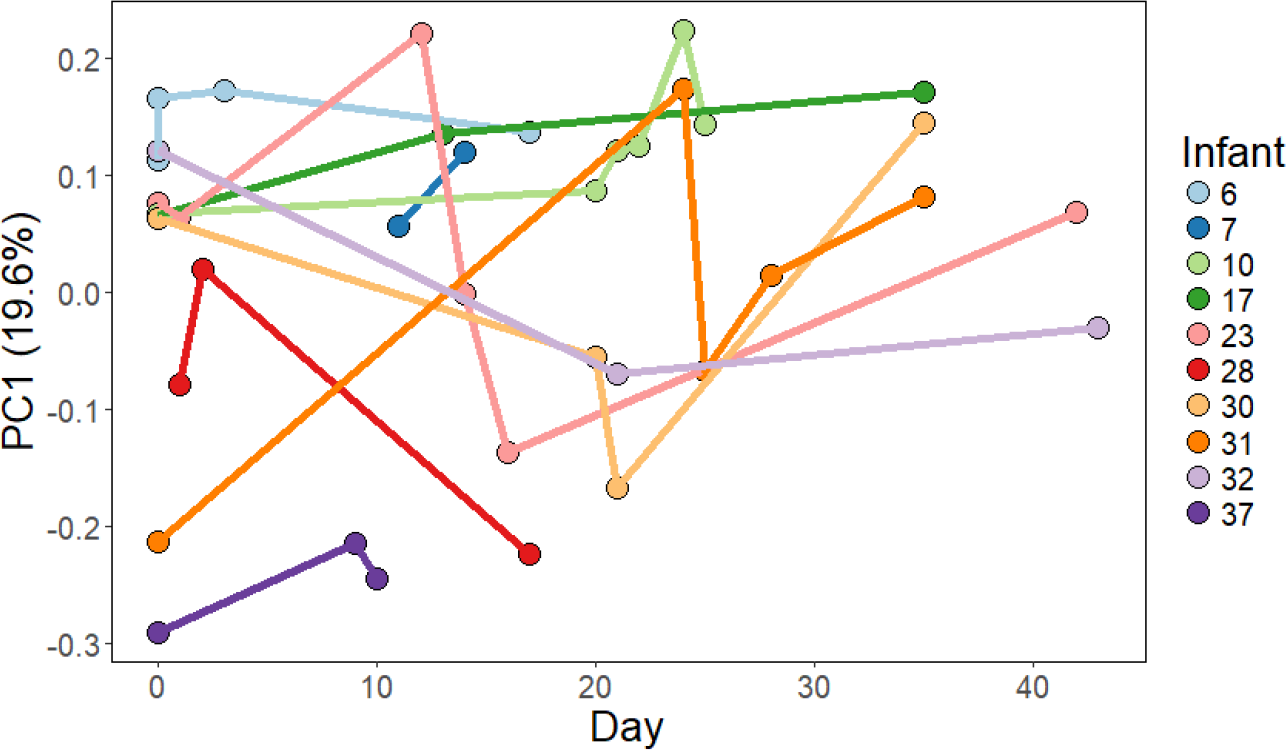
First component of PCoA of metabolite profile over time. Manhattan distances between samples were calculated and visualized by PCoA. The first principal component which explains the most variation among the samples is shown over time. Each dot represents a single fecal sample and is colored by infant. Lines connect samples for each infant to show change over time. Only infants with three or more longitudinal samples shown.

To determine which metabolites might be useful for tracking bacterial metabolism in the infant gut, we examined metabolites with consistent abundance among infants versus those that varied (**Supplemental figure 3**). In general, sugars, central metabolites, and amino acids were variable while fatty acids, sterols, organic acids, and bile acids were more consistent. Infant 23, which developed necrotizing enterocolitis at day 16 of life, had low abundances of amino acid metabolites the two days prior to disease onset (Figure 4). However, several of the healthy control infants also had similarly low abundances of amino acid metabolites. The individual signal of each infant’s metabolome is far more evident than any trends due to clinical factors (**Supplemental table 1**).

### Bacterial composition associated with metabolite profile

Bacterial metabolism in the gut is expected to contribute to the abundances of metabolites detected in fecal samples. We wanted to know if fecal samples with a similar bacterial composition were also similar in their metabolite profile. We employed a Mantel test using Pearson correlations between distances among bacterial compositions of samples and distances among metabolite profiles of samples. Because bacterial compositions and metabolite profiles are personalized, using multiple samples from a single infant would skew the result. Therefore, one sample from each infant was randomly selected 100 times to remove the effect of the individual and the Mantel test was applied to each subset. The average Mantel statistic of r = 0.23 ± 0.05 (p = 0.03 ± 0.03) indicates a weak but significant association between the bacterial composition and metabolite profile. Also, within individual infants, shifts in the bacterial composition are accompanied by shifts in the metabolome. Infants 17, 23, and 31 have dramatic shifts in both bacterial composition and metabolome profile over time, while infants 10 and 37 remain stable in both bacterial composition and metabolome.

To investigate the correlations driving this overall association, we calculated correlations between bacterial abundances and metabolite intensities (Figure 6). *Staphylococcus* had the most positive correlations including several classes of sugar metabolites, organic acids, and central metabolites. Fatty acids, lipid metabolism, and amino acids were positively correlated with the commonly abundant gut colonizers *Enterobacteriaceae* and *Bacteroides*, and negatively correlated with the commonly low abundance colonizers *Staphylococcus* and *Enterococcus*. We also looked more specifically at individual metabolites correlated with bacterial abundances (Figure 6). Bacteroidetes were found to be positively correlated with succinate (r = 0.85). Many other weak correlations (r < 0.5) exist between bacterial abundances and metabolite intensities, but the sample size is not large enough to distinguish signal from noise.

**Figure 6.**
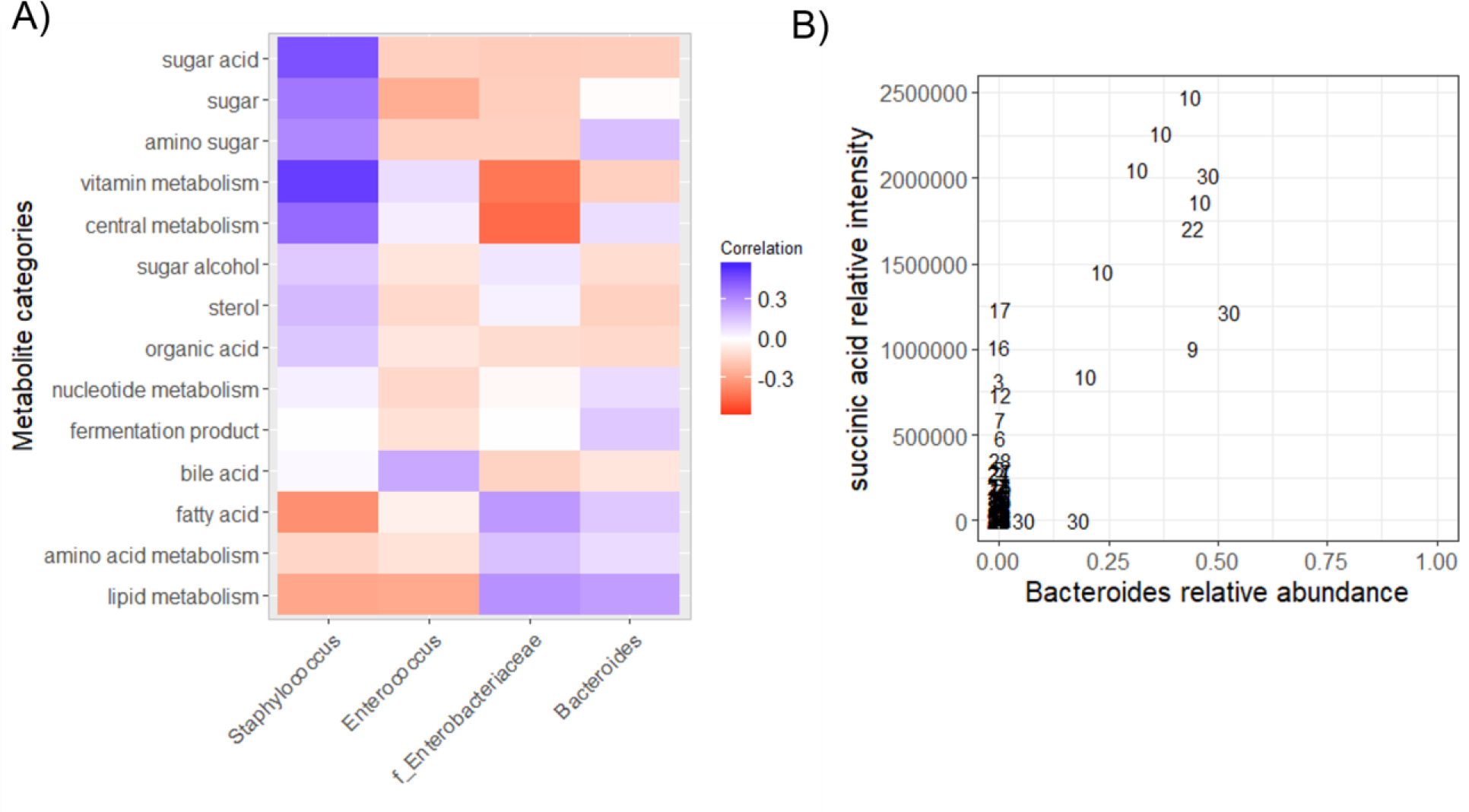
Correlations between bacteria abundances and metabolite intensities. A) Average of correlations between bacterial abundances and all metabolites in each metabolite category. B) Correlation between *Bacteroides* abundance and succinic acid intensity in all samples. Numbers indicate infant number.

## Discussion

Bacterial compositions in this cohort were consistent with the emerging picture from other studies that show the preterm infant gut harbors communities dominated by facultative anaerobes including *Enterobacteriaceae*, *Enterococcus*, and *Staphylococcus* (1, 2, 20). These communities appear to be enriched in commonly antibiotic resistant organisms (21). While we expected to find associations between bacterial community composition and health outcome in this cohort, we were surprised to find that there were not clear signatures of microbiome composition linked to NEC or sepsis. In larger cohorts, associations between particular bacteria or metabolites with NEC have been reported, however, they are not universal signatures across patients, and may reflect patient variation more than disease etiology (22–25). In fact, the strongest signal in both the microbiome and metabolome data from this cohort was the infant from whom the sample was taken. Overall, preterm infant microbiomes in this study were shaped by antibiotics, which have a strong impact on all patients, regardless of health outcome.

Although the bacterial composition of infant guts varied over time, we saw longitudinal samples from individual infants remained highly personalized over several weeks; nearly half of the variation in the microbial community compositions can be explained by which individual the sample came from. The stability of animal-associated microbiomes is an active area of research, with studies finding that the individual microbiome of an adult remains stable through time (26), but can be perturbed by extreme changes in diet or antibiotics (27–29). The bacterial composition in the adult gut largely returns to its previous state one month after antibiotic treatment, but altering the initial assembly of the microbiota in infants can have long lasting health consequences (7, 27, 30, 31). Previous work has found ampicillin and gentamycin (the most common antibiotics taken by infants in this study) to have an inconsistent effect on bacterial diversity, sometimes increasing and sometimes decreasing diversity (1). Similarly, in these infants, ampicillin and gentamycin resulted in more variation in bacterial diversity, but there was no clear trend of increasing or decreasing diversity. However, antibiotics change the dominant members of the microbiota which could have profound effects on immune development and growth (7, 31–33).

Evidence is emerging that a healthy infant gut microbiota is dominated by *Bifidobacteria* with the capacity to digest human milk oligosaccharides in breastmilk (5, 34, 35). The lack of a core *Bifidobacteria* community in infants could leave the microbiota open to colonization by facultative anaerobes like we observed in these infants (36). Proteobacteria such as *Enterobacteriaceae* are commonly seen to increase in abundance after antibiotic administration (21). Indeed, infants in this study had microbiomes that were shaped by antibiotic use. Although six of the thirty-two infants in this study did not have recorded antibiotic use around sampling time, the microbiota can still be affected by prenatal antibiotics taken by the mother (7, 31, 37).

Microbiome studies have become widespread, so that a typical bacterial composition is well characterized in a range of sample cohorts. However, the same cannot be said for the metabolome. There is a dearth of knowledge about what a consensus healthy infant fecal metabolome should be, making comparisons for the cohort in this study difficult. To add to the complexity, each metabolomic approach detects subsets of metabolites, and depends on sample extraction and other method choices. Increasing the frequency of metabolomics data collection in microbiome studies would be a huge step forward for the field. Baseline knowledge about the typical connections between metabolite abundances and bacterial metabolism should be collected, so that molecules that have consistent abundances in a healthy state could give context to data generated from clinical samples in different disease states.

Untargeted metabolomics can survey many metabolites in a biological sample to provide a snapshot of the active metabolism in a system such as the human gut. Metabolite profiles of preterm infants in this study were found to be personalized to a similar degree as the bacterial composition. This is in contrast to a previous study on full term infants that showed the metabolomic profile to be stable, and weakly associated with bacterial composition, over the first few years of life (38). Personalized metabolic signatures of disease hold great promise to complement microbiota profiling in human systems (18, 36). However, analyzing metabolomic data is challenging because an array of inputs contribute to the abundances of metabolites in fecal samples including bacterial metabolism, host biology, and food intake.

We report a number of correlations between bacteria and metabolites in preterm infant feces, and bacterial metabolism has been previously shown to contribute to metabolite abundances in humans and mice (15, 39, 40). Short chain fatty acids are now commonly measured and associated with bacterial fermentation in the gut (41). In this study, the only short-chain fatty acid detected was succinate, which we found to be correlated with the presence of *Bacteroides*, which produces acetate and succinate from carbohydrate fermentation (42). We also detected several medium-chain fatty acids, which were generally correlated with the abundance of *Bacteroides* and *Enterobacteriaceae*. None of the twenty-two C-section born infants in this study were colonized by *Bacteroides*, possibly due to a lack of vertical transmission from the mother during birth (3).

Overall, we find preterm infant microbiomes are shaped by shared exposures especially to antibiotics, leading to the dominance of antibiotic resistant facultative anaerobes such as *Enterococcus* spp.. The anaerobic, milk degrading *Bifidobacteria* were largely absent, even in preterm infants with access to breastmilk, possibly due to a lack of exposure to microbes from family members in the sterile hospital environment along with antibiotics. Our understanding of the health consequences of microbial colonization under these antibiotic-enriched circumstances is still in its infancy.

## Materials and methods

### Sample Collection

Stool samples from diapers of preterm infants in the neonatal intensive care unit at Children’s Hospital Orange County were collected by nurses over three years from 2011 to 2014. Samples were immediately stored at −20 °C then transferred to −80 °C no more than three days post-collection. Samples were kept at −80 °C and thawed once for DNA extraction and metabolomics. A total of 77 stool samples were collected from 32 preterm infants.

### DNA extraction and 16S rRNA gene sequencing

Stool samples were thawed once and DNA was extracted from 10 mg using a fecal Zymo Fecal DNA MiniPrep Kit (#D6010). The V3 and V4 region of the 16SrRNA gene was amplified with two-stage PCR. The first PCR amplified the V3 to V4 region of the 16S rRNA gene using 341F and 805R primers: forward primer (5’- CCTACGGGNGGCWGCAG-3’) and reverse primer (5’- GACTACHVGGGTATCTAATCC −3’) (43). These primers also added an overhang so that barcodes and Illumina adaptors could be added in the second PCR. The first PCR was done as follows: 30 cycles of 95 °C 30 seconds; 65 °C 40 seconds; 72 °C 1 minute. Immediately after completion of the first PCR, primers with sample specific barcodes and Illumina adapter sequences were added and a second PCR was performed as follows: 9 cycles 94 °C for 30 seconds; 55 °C 40 seconds; 72 °C 1 minute. PCR reactions were cleaned using Agencourt AMPure XP magnetic beads (#A63880) using the recommended protocol. Amplicons were run on an agarose gel to confirm amplification and then pooled. Amplicon pool was run on an agarose gel and the 500bp fragment was cut out and gel extracted using Millipore Gel Extraction Kit (#LSKGEL050). The sequencing library was quantified using Quant-iT Pico Green dsDNA Reagent and sent to Laragen Inc. for sequencing on the Illumina MiSeq platform with 250bp paired-end reads producing a total of 2.4 million paired-end reads.

### qPCR for bacterial load

The bacterial load of each fecal sample was measured with quantitative PCR of a conserved region of the 16S gene. The following primers were used: (5’- TCC TAC GGG AGG CAG CAG T-3’), (5’- GGA CTA CCA GGG TAT CTA ATC CTG TT-3’). PerfeCTa SYBER Green SuperMix Reaction Mix (Quantabio #95054) was used to quantify DNA from samples. Relative abundance of 16S rRNA genes relative to the mass of stool was compared for each sample. Total fecal DNA was measured with Quant-iT Pico Green dsDNA Assay Kit (ThermoFisher #P11496).

### Sequence processing

Sequences were quality filtered using PrinSeq to remove adapters as well as any sequences less than 120 base-pairs, containing any ambiguous bases, or with a mean PHRED quality score of less than 30 (44). Reads were found to drop steeply in quality after 140 base pairs, so all reads were trimmed to 140 base pairs. The forward read contained the V3 region in the high quality first 140 base pairs, while the V4 region was captured in the low-quality region of the reverse reads. Therefore, we used only the forward reads for subsequent analyses.

### Bacterial community analysis

Quantitative Insights Into microbial Ecology (QIIME) was used for de novo OTUs picking using the Swarm algorithm with a clustering threshold of 8 (45, 46). This resulted in 2,810 OTUs among all samples. OTUs containing only one sequence were filtered out, leaving 212 OTUs. Taxonomy was assigned to each OTU using QIIME and the Greengenes 13_8 database. An OTU table was constructed and used for downstream analysis. Ten rarefactions were performed on the OTU table down to 2000 reads per sample, which was the largest number of reads that allowed retention of most samples. QIIME was used to calculate alpha diversity by Shannon index and beta diversity by the average weighted UniFrac distance of the ten rarefactions. Community composition barplots, Principal Coordinate Analysis (PCoA) plots, and alpha diversity plots were created using R and the ggplot2 package (47, 48). All R scripts are included in the supplemental information.

### Untargeted metabolomics by GC-MS

When fecal samples were thawed for DNA extraction, approximately 50 mg was collected and refrozen at −80 ° for metabolomics. Samples were sent on dry ice to the West Coast Metabolomics Center (WCMC) at UC Davis for untargeted metabolomics by gas chromatography time-of-flight mass spectrometry. Metabolites were extracted from fecal samples with a 3:3:2 mixture of isopropanol, acetonitrile, and water respectively before derivatization and GC-MS analysis by Fiehn Lab standard operating procedures (49–51). Metabolite profiles were compared by calculating Manhattan distances between samples based on all metabolite intensities and visualized by PCoA using the vegan and ape packages in R (52, 53).

### Permutational multivariate analysis of variance (PERMANOVA)

PERMANOVA was used to determine factors that explained variance in bacterial community composition and metabolite profile. PERMANOVA was performed using the adonis function in the vegan package in R. The input for PERMANOVA was UniFrac distances of the 16S data and Manhattan distances of the metabolite profiles. Briefly, PERMANOVA quantifies the variation among samples explained by the given groupings compared to randomized groupings. To measure the variance explained by individual infant, we excluded samples that had fewer than three longitudinal samples, leaving ten infants. When performing PERMANOVA for factors other than individual, we accounted for the longitudinal sampling by repeatedly subsampling one sample from each infant and averaging the results.

### Correlations between bacteria and metabolites

Pearson correlations between bacterial abundances and normalized metabolite intensities were calculated using the cor function in R. Correlations were calculated between the relative abundances of all bacterial classes and all metabolite intensities among all samples in all infants. Only the four most highly abundant general of bacteria were used to ensure no results were skewed by taxa present in only one or a few samples. For each class of metabolite, the average of all correlations between metabolites in that class and each taxon was calculated so that trends between bacterial taxa and classes of metabolites could be visualized by heatmap.

### Mantel test

To determine if fecal samples with similar bacterial compositions also have similar metabolite profiles, a Mantel test was performed. To account for the effect of longitudinal sampling, each dataset was randomly subsampled down to one sample per infant. A Bray-Curtis dissimilarity matrix was computed for the bacterial composition and Manhattan distances calculated for metabolite intensities. The Mantel function in the vegan package of R was used to calculate the Mantel statistic for a Pearson correlation between the two dissimilarity matrices. The average and standard deviation of the Mantel statistic r and p-value for the 100 Mantel tests was reported.

## Data availability

Raw sequence data will be uploaded to the SRA. OTU tables, raw metabolomics data, a markdown file of sequence processing workflow, and R scripts used for analyses are available at https://github.com/swandro/preterm_infant_analysis.

## Acknowledgements

This project was supported by a UCI Single Investigator grant, the CORCCL SIIG-2014-2015-51, a pilot grant from the UC Davis West Coast Metabolomics Core as part of NIH-DK097154, and start-up funds for the Whiteson Lab in UC Irvine’s Molecular Biology and Biochemistry Department. Thanks to CORCL for funding.

Thank you to Ying (Lucy) Lu for help with qPCR experiments. Thank you Celine Mouginot and Adam Martiny for providing 16S primers and protocols. Thanks to Ilhem Messaoudi and members of the Whiteson lab for editing. Thanks to Megan Showalter and Prof. Oliver Fiehn of the UC Davis West Coast Metabolomics Core were very helpful in carrying out untargeted GC-MS profiling.

## References

1. Gibson MK, Wang B, Ahmadi S, Burnham C-AD, Tarr PI, Warner BB, Dantas G. 2016. Developmental dynamics of the preterm infant gut microbiota and antibiotic resistome. Nat Microbiol 1:16024.

2. Rosa PS La, Warner BB, Zhou Y, Weinstock GM, Sodergren E, Hall-Moore CM, Stevens HJ, Bennett WE, Shaikh N, Linneman LA, Hoffmann JA, Hamvas A, Deych E, Shands BA, Shannon WD, Tarr PI. 2014. Patterned progression of bacterial populations in the premature infant gut. Proc Natl Acad Sci 111: 12522–12527.

3. Bäckhed F, Roswall J, Peng Y, Feng Q, Jia H, Kovatcheva-Datchary P, Li Y, Xia Y, Xie H, Zhong H, Khan MT, Zhang J, Li J, Xiao L, Al-Aama J, Zhang D, Lee YS, Kotowska D, Colding C, Tremaroli V, Yin Y, Bergman S, Xu X, Madsen L, Kristiansen K, Dahlgren J, Wang J. 2015. Dynamics and Stabilization of the Human Gut Microbiome during the First Year of Life. Cell Host Microbe 17: 690–703.

4. Bokulich NA, Chung J, Battaglia T, Henderson N, Jay M, Li H, D Lieber A, Wu F, Perez-Perez GI, Chen Y, Schweizer W, Zheng X, Contreras M, Dominguez-Bello MG, Blaser MJ. 2016. Antibiotics, birth mode, and diet shape microbiome maturation during early life. Sci Transl Med 8:343ra82.

5. Frese SA, Hutton AA, Contreras LN, Shaw CA, Palumbo MC, Casaburi G, Xu G, Davis JCC, Lebrilla CB, Henrick BM, Freeman SL, Barile D, German JB, Mills DA, Smilowitz JT, Underwood MA. 2017. Persistence of Supplemented Bifidobacterium longum subsp. infantis EVC001 in Breastfed Infants. mSphere 2: e00501-17.

6. Nobel YR, Cox LM, Kirigin FF, Bokulich NA, Yamanishi S, Teitler I, Chung J, Sohn J, Barber CM, Goldfarb DS, Raju K, Abubucker S, Zhou Y, Ruiz VE, Li H, Mitreva M, Alekseyenko A V, Weinstock GM, Sodergren E, Blaser MJ. 2015. Metabolic and metagenomic outcomes from early-life pulsed antibiotic treatment. Nat Commun 6:7486.

7. Cox LM, Yamanishi S, Sohn J, Alekseyenko A V, Leung JM, Cho I, Kim SG, Li H, Gao Z, Mahana D, Zárate Rodriguez JG, Rogers AB, Robine N, Loke P, Blaser MJ. 2014. Altering the Intestinal Microbiota during a Critical Developmental Window Has Lasting Metabolic Consequences. Cell 158: 705–721.

8. Tanaka S, Kobayashi T, Songjinda P, Tateyama A, Tsubouchi M, Kiyohara C, Shirakawa T, Sonomoto K, Nakayama J. 2009. Influence of antibiotic exposure in the early postnatal period on the development of intestinal microbiota. FEMS Immunol Med Microbiol 56: 80–87.

9. Keski-Nisula L, Kyynäräinen H-R, Kärkkäinen U, Karhukorpi J, Heinonen S, Pekkanen J. 2013. Maternal intrapartum antibiotics and decreased vertical transmission of Lactobacillus to neonates during birth. Acta Paediatr (Oslo, Norw 1992) 102: 480–485.

10. Ting JY, Synnes A, Roberts A, Deshpandey A, Dow K, Yoon EW, Lee K-S, Dobson S, Lee SK, Shah PS. 2016. Association Between Antibiotic Use and Neonatal Mortality and Morbidities in Very Low-Birth-Weight Infants Without Culture-Proven Sepsis or Necrotizing Enterocolitis. JAMA Pediatr 170: 1181–1187.

11. Stokholm J, Schjørring S, Pedersen L, Bischoff AL, Følsgaard N, Carson CG, Chawes BLK, Bønnelykke K, Mølgaard A, Krogfelt KA, Bisgaard H. 2013. Prevalence and predictors of antibiotic administration during pregnancy and birth. PLoS One 8:e82932.

12. Fitzgibbons SC, Ching Y, Yu D, Carpenter J, Kenny M, Weldon C, Lillehei C, Valim C, Horbar JD, Jaksic T. 2009. Mortality of necrotizing enterocolitis expressed by birth weight categories. J Pediatr Surg 44: 1072–1076.

13. Nanthakumar N, Meng D, Goldstein AM, Zhu W, Lu L, Uauy R, Llanos A, Claud EC, Walker WA. 2011. The Mechanism of Excessive Intestinal Inflammation in Necrotizing Enterocolitis: An Immature Innate Immune Response. PLoS One 6: e17776.

14. Dodd D, Spitzer MH, Van Treuren W, Merrill BD, Hryckowian AJ, Higginbottom SK, Le A, Cowan TM, Nolan GP, Fischbach MA, Sonnenburg JL. 2017. A gut bacterial pathway metabolizes aromatic amino acids into nine circulating metabolites. Nature 551:648.

15. Wikoff WR, Anfora AT, Liu J, Schultz PG, Lesley SA, Peters EC, Siuzdak G. 2009. Metabolomics analysis reveals large effects of gut microflora on mammalian blood metabolites. Proc Natl Acad Sci U S A 106: 3698–703.

16. Willemsen LEM, Koetsier MA, van Deventer SJH, van Tol EAF. 2003. Short chain fatty acids stimulate epithelial mucin 2 expression through differential effects on prostaglandin E1 and E2 production by intestinal myofibroblasts. Gut 52: 1442–1447.

17. Chang P V, Hao L, Offermanns S, Medzhitov R. 2014. The microbial metabolite butyrate regulates intestinal macrophage function via histone deacetylase inhibition. Proc Natl Acad Sci 111: 2247–2252.

18. Kelly CJ, Zheng L, Campbell EL, Saeedi B, Scholz CC, Bayless AJ, Wilson KE, Glover LE, Kominsky DJ, Magnuson A, Weir TL, Ehrentraut SF, Pickel C, Kuhn KA, Lanis JM, Nguyen V, Taylor CT, Colgan SP. 2015. Crosstalk between Microbiota-Derived Short-Chain Fatty Acids and Intestinal Epithelial HIF Augments Tissue Barrier Function. Cell Host Microbe 17: 662–671.

19. Cole JR, Wang Q, Fish JA, Chai B, McGarrell DM, Sun Y, Brown CT, Porras-Alfaro A, Kuske CR, Tiedje JM. 2014. Ribosomal Database Project: data and tools for high throughput rRNA analysis. Nucleic Acids Res 42:D633-42.

20. Grier A, Qiu X, Bandyopadhyay S, Holden-Wiltse J, Kessler HA, Gill AL, Hamilton B, Huyck H, Misra S, Mariani TJ, Ryan RM, Scholer L, Scheible KM, Lee Y-H, Caserta MT, Pryhuber GS, Gill SR. 2017. Impact of prematurity and nutrition on the developing gut microbiome and preterm infant growth. Microbiome 5:158.

21. Sommer MO, Dantas G. 2011. Antibiotics and the resistant microbiome. Curr Opin Microbiol 14: 556–563.

22. Morrow AL, Lagomarcino AJ, Schibler KR, Taft DH, Yu Z, Wang B, Altaye M, Wagner M, Gevers D, Ward D V, Kennedy MA, Huttenhower C, Newburg DS. 2013. Early microbial and metabolomic signatures predict later onset of necrotizing enterocolitis in preterm infants. Microbiome 1:13.

23. Sim K, Shaw AG, Randell P, Cox MJ, McClure ZE, Li M-S, Haddad M, Langford PR, Cookson WOCM, Moffatt MF, Kroll JS. 2015. Dysbiosis Anticipating Necrotizing Enterocolitis in Very Premature Infants. Clin Infect Dis 60: 389–397.

24. Heida FH, Zoonen V, F AGJ, Hulscher JBF, Kiefte T, C BJ, Wessels R, Kooi EMW, Bos AF, Harmsen HJM, Goffau D, C M. A Necrotizing Enterocolitis-Associated Gut Microbiota Is Present in the Meconium: Results of a Prospective Study 62: 863–870.

25. Cassir N, Benamar S, Khalil JB, Croce O, Saint-Faust M, Jacquot A, Million M, Azza S, Armstrong N, Henry M, Jardot P, Robert C, Gire C, Lagier J-C, Chabrière E, Ghigo E, Marchandin H, Sartor C, Boutte P, Cambonie G, Simeoni U, Raoult D, Scola B La. Clostridium butyricum Strains and Dysbiosis Linked to Necrotizing Enterocolitis in Preterm Neonates 61: 1107–1115.

26. Faith JJ, Guruge JL, Charbonneau M, Subramanian S, Seedorf H, Goodman AL, Clemente JC, Knight R, Heath AC, Leibel RL, Rosenbaum M, Gordon JI. 2013. The Long-Term Stability of the Human Gut Microbiota. Science (80-) 341: 1237439–1237439.

27. Dethlefsen L, Relman DA. 2011. Incomplete recovery and individualized responses of the human distal gut microbiota to repeated antibiotic perturbation. Proc Natl Acad Sci U S A 108 Suppl: 4554–4561.

28. David LA, Maurice CF, Carmody RN, Gootenberg DB, Button JE, Wolfe BE, Ling A V., Devlin AS, Varma Y, Fischbach MA, Biddinger SB, Dutton RJ, Turnbaugh PJ. 2013. Diet rapidly and reproducibly alters the human gut microbiome. Nature 505: 559–563.

29. Brüssow H. 2016. How stable is the human gut microbiota? And why this question matters. Environ Microbiol 18: 2779–2783.

30. Dethlefsen L, Huse S, Sogin ML, Relman DA. 2008. The Pervasive Effects of an Antibiotic on the Human Gut Microbiota, as Revealed by Deep 16S rRNA Sequencing. PLOS Biol 6:e280.

31. Schulfer A, Blaser MJ. 2015. Risks of Antibiotic Exposures Early in Life on the Developing Microbiome. PLOS Pathog 11:e1004903.

32. Metsälä J, Lundqvist A, Virta LJ, Kaila M, Gissler M, Virtanen SM. 2015. Prenatal and post-natal exposure to antibiotics and risk of asthma in childhood. Clin Exp Allergy 45: 137–145.

33. Mueller NT, Whyatt R, Hoepner L, Oberfield S, Dominguez-Bello MG, Widen EM, Hassoun A, Perera F, Rundle A. 2015. Prenatal exposure to antibiotics, cesarean section and risk of childhood obesity. Int J Obes 39: 665–670.

34. Karav S, Parc A Le, Bell JMLN de M, Frese SA, Kirmiz N, Block DE, Barile D, Mills DA. 2016. Oligosaccharides Released from Milk Glycoproteins Are Selective Growth Substrates for Infant-Associated Bifidobacteria.

35. Underwood MA, Davis JCC, Kalanetra KM, Gehlot S, Patole S, Tancredi DJ, Mills DA, Lebrilla CB, Simmer K. 2017. Digestion of Human Milk Oligosaccharides by Bifidobacterium Breve in the Premature Infant. J Pediatr Gastroenterol Nutr 1.

36. Stewart CJ, Embleton ND, Marrs ECL, Smith DP, Nelson A, Abdulkadir B, Skeath T, Petrosino JF, Perry JD, Berrington JE, Cummings SP. 2016. Temporal bacterial and metabolic development of the preterm gut reveals specific signatures in health and disease. Microbiome 4.

37. Clark RH, Bloom BT, Spitzer AR, Gerstmann DR. 2006. Reported Medication Use in the Neonatal Intensive Care Unit: Data From a Large National Data Set. Pediatrics 117: 1979–1987.

38. Kostic AD, Gevers D, Siljander H, Vatanen T, Hyötyläinen T, Hämäläinen A-M, Peet A, Tillmann V, Pöhö P, Mattila I, Lähdesmäki H, Franzosa EA, Vaarala O, de Goffau M, Harmsen H, Ilonen J, Virtanen SM, Clish CB, Orešič M, Huttenhower C, Knip M,DIABIMMUNE Study Group RJ, Xavier RJ. 2015. The dynamics of the human infant gut microbiome in development and in progression toward type 1 diabetes. Cell Host Microbe 17: 260–73.

39. Yen S, McDonald JAK, Schroeter K, Oliphant K, Sokolenko S, Blondeel EJM, Allen-Vercoe E, Aucoin MG. 2015. Metabolomic Analysis of Human Fecal Microbiota: A Comparison of Feces-Derived Communities and Defined Mixed Communities. J Proteome Res 14: 1472–1482.

40. Dodd D, Spitzer MH, Van Treuren W, Merrill BD, Hryckowian AJ, Higginbottom SK, Le A, Cowan TM, Nolan GP, Fischbach MA, Sonnenburg JL. 2017. A gut bacterial pathway metabolizes aromatic amino acids into nine circulating metabolites. Nature 551:648.

41. Morrison DJ, Preston T. 2016. Formation of short chain fatty acids by the gut microbiota and their impact on human metabolism. Gut Microbes 7: 189–200.

42. Miller TL. 1978. The pathway of formation of acetate and succinate from pyruvate by Bacteroides succinogenes. Arch Microbiol 117: 145–52.

43. Herlemann DP, Labrenz M, Jürgens K, Bertilsson S, Waniek JJ, Andersson AF. 2011. Transitions in bacterial communities along the 2000 km salinity gradient of the Baltic Sea. ISME J 5: 1571–1579.

44. Schmieder R, Edwards R. 2011. Quality control and preprocessing of metagenomic datasets. Bioinformatics 27: 863–864.

45. Caporaso JG, Kuczynski J, Stombaugh J, Bittinger K, Bushman FD, Costello EK, Fierer N, Peña AG, Goodrich JK, Gordon JI, Huttley GA, Kelley ST, Knights D, Koenig JE, Ley RE, Lozupone CA, McDonald D, Muegge BD, Pirrung M, Reeder J, Sevinsky JR, Turnbaugh PJ, Walters WA, Widmann J, Yatsunenko T, Zaneveld J, Knight R. 2010. QIIME allows analysis of high-throughput community sequencing data. Nat Methods 7: 335–336.

46. Mahé F, Rognes T, Quince C, de Vargas C, Dunthorn M. 2014. Swarm: robust and fast clustering method for amplicon-based studies. PeerJ 2:e593.

47. Wickham H. 2009. ggplot2: Elegant Graphics for Data Analysis. Springer-Verlag New York.

48. Lozupone C, Knight R. 2005. UniFrac: a new phylogenetic method for comparing microbial communities. Appl Environ Microbiol 71: 8228–35.

49. Kind T, Wohlgemuth G, Lee DY, Lu Y, Palazoglu M, Shahbaz S, Fiehn O. 2009. FiehnLib: Mass Spectral and Retention Index Libraries for Metabolomics Based on Quadrupole and Time-of-Flight Gas Chromatography/Mass Spectrometry. Anal Chem 81: 10038–10048.

50. Kind T, Tsugawa H, Cajka T, Ma Y, Lai Z, Mehta SS, Wohlgemuth G, Barupal DK, Showalter MR, Arita M, Fiehn O. 2017. Identification of small molecules using accurate mass MS/MS search. Mass Spectrom Rev.

51. Cajka T, Fiehn O. 2016. Toward Merging Untargeted and Targeted Methods in Mass Spectrometry-Based Metabolomics and Lipidomics. Anal Chem 88: 524–545.

52. Paradis E, Claude J, Strimmer K. 2004. A{PE}: analyses of phylogenetics and evolution in {R} language. Bioinformatics 20: 289–290.

53. Oksanen J, Blanchet FG, Friendly M, Kindt R, Legendre P, McGlinn D, Minchin PR, O’Hara RB, Simpson GL, Solymos P, Stevens MHH, Szoecs E, Wagner H. 2016. vegan: Community Ecology Package.

